# Allosteric covalent inhibition of cereblon through cryptic cysteine targeting

**DOI:** 10.64898/2026.10.10.758245

**Authors:** Zeba Rizvi, Brian E. Nordin, Kent T. Symons, Kristen A Baltgalvis, David S. Weinstein, Matthew P. Patricelli, Gabriel M. Simon, Gabriel C. Lander

**Affiliations:** Department of Integrative Structural and Computational Biology, Scripps Research, La Jolla, California 92307, United States; Vividion Therapeutics, San Diego, California 92121, United States

**Keywords:** CRBN E3 Ligase, Covalent cysteine profiling, VVD-105917 inhibitor, cryo-EM, conformational heterogeneity, sign switchable allosteric modulation, UPS therapeutics

## Abstract

Thalidomide-derived cereblon (CRBN) modulators are among the most clinically impactful targeted protein degradation therapeutics, redirecting the CRL4-CRBN E3 ubiquitin ligase to degrade disease-driving neosubstrates. All current CRBN-targeting IMiDs and CELMoDs engage a single orthosteric pocket, leaving CRBN’s conformational landscape largely unexploited pharmacologically. Using cysteine-directed chemoproteomic screening, we identified covalent ligands that engage Cys287, a cryptic allosteric pocket distal to the orthosteric site. Our lead compound, VVD-105917, acts as a pan-inhibitor of IMiD/CELMoD-driven neosubstrate degradation. Cryo-EM structures reveal that covalent Cys287 engagement traps CRBN in its open, inactive conformation, resolving a previously unobserved N-terminal segment and sensor loop. Sequential incubation experiments show the conformational outcome depends on ligand addition order, demonstrating that even potent CELMoD-driven closure can be overridden by allosteric engagement. A recently reported ligand at this same pocket produces the opposite effect, promoting rather than blocking closure, indicating the Cys287 site is mechanistically sign-switchable. These findings establish covalent chemoproteomics as a strategy for discovering and controlling E3 ligase conformational switches.

## Introduction

The ubiquitin-proteasome system (UPS) orchestrates protein quality control by targeting damaged, misfolded, or regulatory proteins for degradation, thereby ensuring cellular homeostasis ^1^. E3 ubiquitin ligases confer substrate specificity within this system, marking selected proteins for proteasomal clearance ^2^. Their importance is underscored by the fact that dysregulation of E3 ligases contribute to cancer, neurodegeneration, and immune disorders ^3^.

Cereblon (CRBN), a substrate receptor for the CRL4-CRBN E3 ligase complex, was previously identified as the primary target of thalidomide ^4^, a compound notorious for its teratogenicity in humans. However, its repurposing spawned a new class of immunomodulatory imide drugs (IMiDs), including lenalidomide and pomalidomide, which have transformed cancer therapy. These compounds redirect CRBN to degrade zinc finger transcription factors such as Ikaros (IKZF1) and Aiolos (IKZF3), exerting potent anti-myeloma activity ^5^. Structural studies revealed that IMiDs function as molecular glues, creating neomorphic interfaces that enable CRBN to recruit neosubstrates for ubiquitination ^6–8^. Cryoelectron microscopy (cryo-EM) analyses further demonstrated that IMiD binding drives an open-to-closed conformational transition in CRBN, and that this structural switch is essential for productive neosubstrate engagement ^9,13^.

The clinical impact of IMiDs has been transformative. Lenalidomide and pomalidomide are FDA-approved and widely used in multiple myeloma treatment, while newer analogs including iberdomide, mezigdomide, and avadomide (known as cereblon E3 ligase modulators, or CELMoDs) are actively being pursued in clinical trials ^11^. Although these compounds all engage CRBN in a well-defined orthosteric pocket ^6–8,12^, structural studies have shown that CRBN neo-function is ultimately governed by a dynamic conformational landscape ^9,13,14^. Recent discovery of a conserved, cryptic allosteric pocket further indicates that orthosteric engagement is only one layer of CRBN neofunction regulation, and that allosteric ligands can tune CRBN conformational equilibria to modulate neosubstrate degradation ^13^.

## Results & Discussion

### Chemoproteomic discovery of a covalent allosteric CRBN inhibitor

Chemoproteomic strategies targeting reactive cysteine residues have emerged as powerful tools for uncovering cryptic pockets across the proteome ^15–17^. By measuring changes in cysteine probe-reactivity, covalent engagement and conformational changes can be detected on proteins in their native state and unexpected ligandable hotspots can be identified and exploited for drug- and probe discovery. Using this methodology, we discovered covalent ligands that simultaneously block probe-labeling at two cysteine residues on CRBN: Cys287 and Cys318 (**Fig. 1a,b**, **Supplementary Fig. 1a**). Notably, C287 is in the allosteric pocket identified by Dippon et al., ^13^ although this was unknown to us at the time. Given the tight correlation in labeling profiles across a large range of compounds (**Fig. 1b**), and the fact that Cys287 and Cys318 are not proximal, we suspected covalent engagement of one residue was leading to a conformational change that prevents probe-labeling of the other. To determine which residue was directly engaged we performed experiments in which proteins were denatured following compound-treatment and before probe-labeling. Inhibition of probe-labeling was preserved at Cys287, demonstrating covalent attachment to this cysteine, whereas C318 labeling was restored under denaturing conditions suggesting that a conformational change rather than direct engagement was responsible for the disruption of probe-binding at C318 (**Supplementary Fig. 1a**).

**Figure 1.**
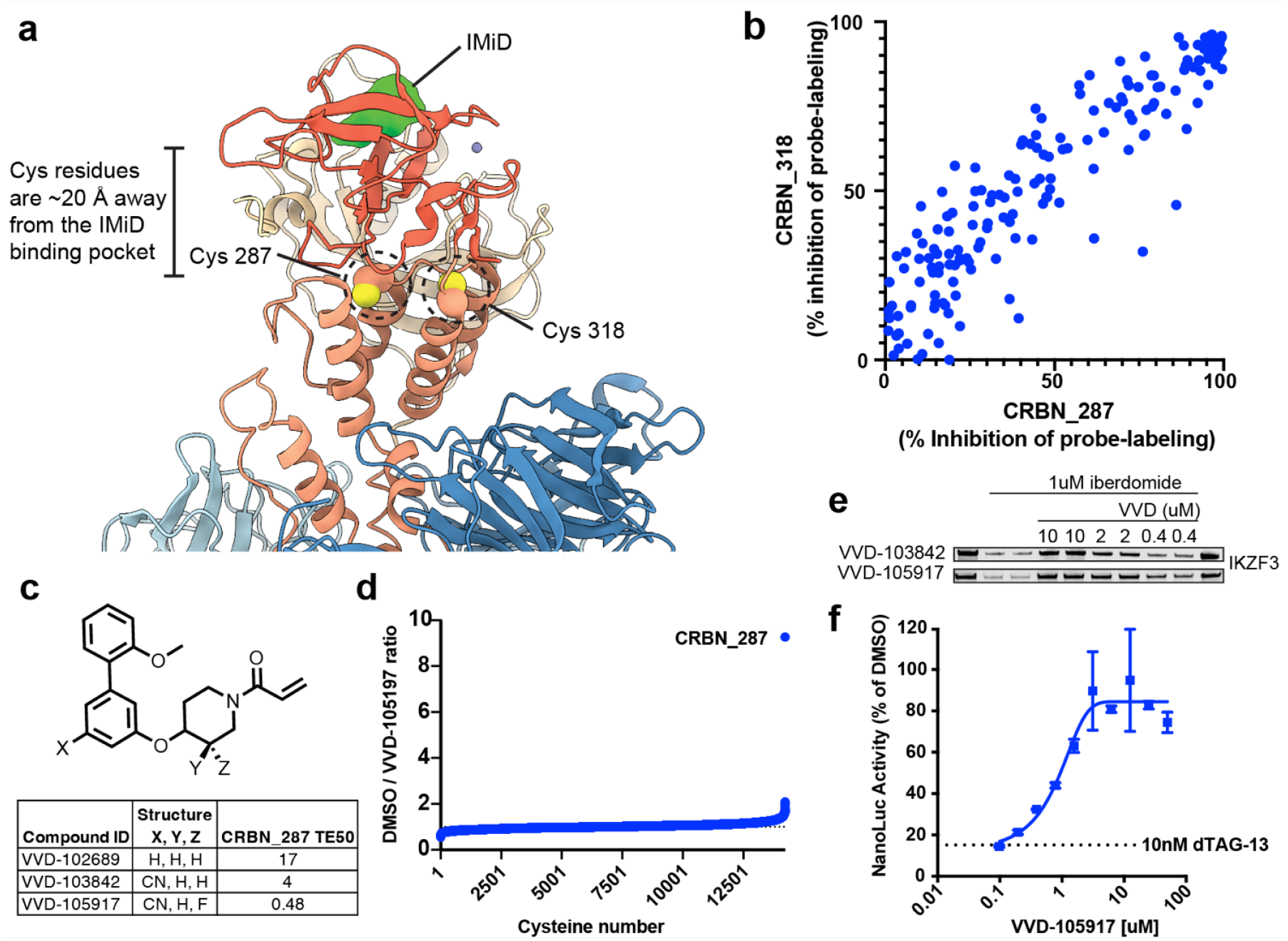
Chemoproteomic discovery and optimization of a covalent CRBN allosteric inhibitor. (**a**). Cryo-EM structure of the closed DDB1^ΔBPB^/CRBN complex (PDB 9OTY). DDB1^ΔBPB^ is shown in blue; CRBN is colored by domain: Lon domain, tan; helical bundle (HB), light red; and thalidomide-binding domain (TBD), red. The immunomodulatory drug (IMiD; lime) occupies the tri-Trp pocket in the TBD. Yellow spheres indicate Cys287 and Cys318, located approximately 20 Å from the canonical IMiD-binding pocket. (**b**). Correlation between inhibition of probe labeling at Cys287 (x axis) and Cys318 (y axis) across a covalent compound library screened at 500 µM by targeted proteomics in Jurkat or PBMC lysates. (**c**). Structure-activity relationships of meta-biphenyl piperazine acrylamide analogs. Substituents at positions X, Y and Z were varied to optimize covalent target engagement. CRBN Cys287 TE50 values were determined by competitive probe labeling. **(d)**. Global cysteine selectivity of VVD-105917 assessed by TMT chemoproteomics in Jurkat cell lysates treated with 10 µM VVD-105917. **(e)**. Dose-dependent inhibition of iberdomide-mediated IKZF3 degradation by VVD-105917 and VVD-103842. MM.1S cells were pretreated with the indicated concentrations of VVD-105917 or VVD-103842 for 2 h and then treated with 1 µM iberdomide for 4 h. IKZF3 (Aiolos) levels were assessed by immunoblotting with an anti-Aiolos antibody; DMSO-treated cells served as the control. **(f)**. Inhibition of dTAG-13-mediated FKBP12 degradation by VVD-105917. HEK293LX cells expressing FKBP12^F36V^-NanoLuc were pretreated with VVD-105917 for 2 h before addition of 10 nM dTAG-13, and FKBP12 levels were assessed after 4 h by HiBiT luminescence.

A series of meta-biphenyl piperazine acrylamides was optimized culminating in VVD-105917, which demonstrated selective and potent covalent engagement with Cys287 (**Fig. 1c,d**). Remarkably, we found that VVD-105917 inhibits iberdomide-mediated IKZF3 degradation in a dose-dependent manner consistent with its TE50, demonstrating that binding to this cryptic pocket inhibits CRBN neofunction (**Fig. 1e**). This inhibitory activity extended to two distinct bifunctional proximity-based degraders: dTAG-13, which targets FKBP12, and dBET1, which degrades BRD4 (**Fig. 1f, Supplementary Fig. 1b**). The consistent inhibition across these platforms demonstrates that VVD-105917 functions as a pan-inhibitor of CRBN-dependent neosubstrate degradation.

### VVD-105917 traps CRBN in an inhibited open conformation

The potent inhibitory activity of VVD-105917 was surprising, given that Cys287 is located 15-25 Å away from the orthosteric tri-Trp thalidomide-binding pocket (TBP). Interaction with a cysteine at this distance would not be expected to directly interfere with IMiD binding, suggesting that VVD-105917 inhibits neofunction through an allosteric mechanism, rather than simple steric occlusion of the TBP. We also discovered that binding of IMiDs partially blocked probe-labeling at C287 and C318 (**Supplementary Fig. 1c**) suggesting that C287 ligands may exclusively bind the open-conformation and that the IMiD-bound closed conformation prevents access to the C287 pocket. To further characterize these findings, we pursued structural studies to elucidate how covalent modification at this distal cysteine could antagonize CRBN neofunction.

A single particle cryo-EM structure of VVD-105917 bound to DDB1^ΔBPB^-CRBN revealed that the ligand binds at the top of the helical bundle domain of CRBN, positioned like a steric wedge that prevents the open-to-close transition of CRBN domains, thereby trapping CRBN in the therapeutically inactive open conformation (PDB:38MG; from this study) (**Fig. 2a**). Notably, in the VVD-105917-bound structure we observe a rotameric rearrangement of Trp264, which is proximal to the Cys287 adduct (**Fig. 2a**). This rearrangement is consistent with an induced accommodation of the ligand that may contribute to stability of the open CRBN conformer.

**Figure 2.**
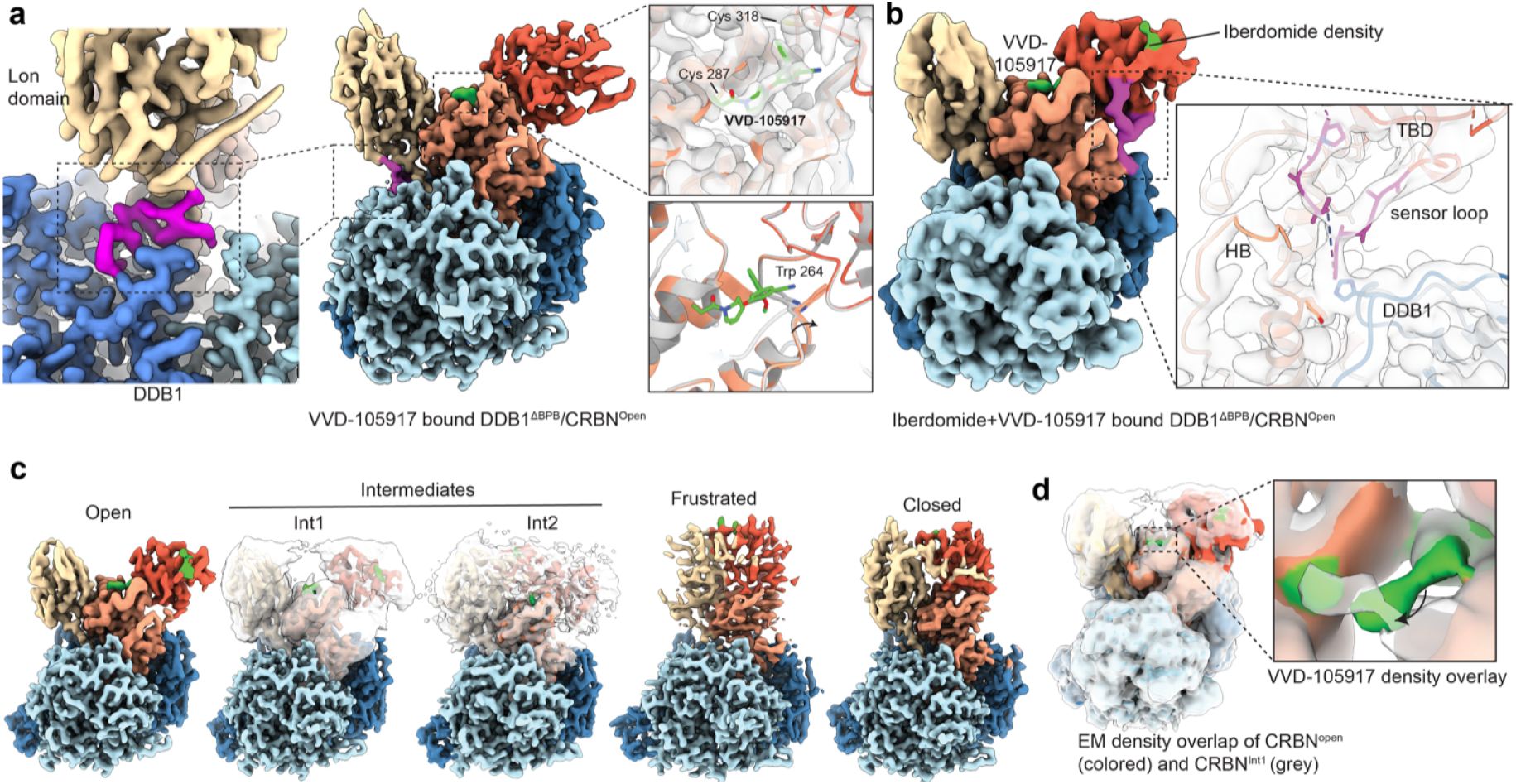
Cryo-EM structure of VVD-105917 bound CRBN reveals an inhibited open conformation. **(a)**, Center, composite cryo-EM map of the DDB1^ΔBPB^/CRBN complex determined in the presence of VVD-105917. The map is colored by component: CRBN Lon domain, tan; thalidomide-binding domain (TBD), red; helical bundle (HB), light red; DDB1^ΔBPB^ β-propellers A and C, light and dark blue, respectively; VVD-105917, green; and putative CRBN N-terminal density, magenta. Left, enlarged view of the putative N-terminal density at the CRBN-DDB1 interface. Right, the upper panel shows VVD-105917 covalently linked to Cys287 in the allosteric pocket; Cys318 remains unmodified. The lower panel shows the rotameric rearrangement of Trp264 that accommodates VVD-105917. **(b)**, Composite cryo-EM map of DDB1^ΔBPB^/CRBN determined in the presence of iberdomide and VVD-105917, colored as in a. Density for iberdomide corresponds to its glutarimide-isoindolinone core. Inset, cryo-EM density and atomic model for the sensor loop (magenta) in the open conformation. (**c**), Representative composite cryo-EM maps of DDB1^ΔBPB^/CRBN in the Open, Intermediate 1 (Int1), Intermediate 2 (Int2), Frustrated, and Closed conformational states, colored as in a. Density for VVD-105917 (green) is visible in the Open and intermediate states.(**d**), Conformational heterogeneity of the allosteric pocket in the Open and Int1 states. Left, overlay of the Open and Int1 cryo-EM maps; the Int1 map was low-pass filtered to emphasize global conformational differences. Right, enlarged view showing that the VVD-105917 density is shifted in Int1 relative to the Open state, consistent with remodeling of the allosteric pocket during CRBN conformational rearrangement.

Unexpectedly, we observe a previously unreported density at the interface between the Lon domain of CRBN and DDB1 (**Fig. 2a, left insets**). Although the density is not sufficiently resolved for unambiguous atomic modeling, we hypothesize that it corresponds to a segment of the CRBN N-terminal region that becomes stabilized in the long-lived open conformation induced by VVD-105917. Two alternate putative models for this segment are presented in (**Supplementary Fig. 2**).

### VVD-105917 remodels CELMoD-bound CRBN conformational ensembles

To define how VVD-105917 influences CELMoD-CRBN conformational landscape, we next conducted sequential incubation experiments with iberdomide or mezigdomide (**Table 1**). We were intrigued by our observation that the order of compound addition influenced the conformational outcomes. When VVD-105917 was added prior to the addition of iberdomide, the entire CRBN population remained in the open conformation, indicating that iberdomide is unable to overcome inhibition by the covalent ligand (**Table 1**). In the reciprocal experiment, CRBN was first incubated with iberdomide for 10 minutes - conditions previously shown to close approximately 50% of the CRBN population ^9^ - followed by the addition of VVD-105917 for 30 minutes. Strikingly, only 9.5% were found in closed conformation, whereas 52% of CRBN molecules remained in the open state, and the remaining 38.5% was found to be shifted in an “intermediate” state, where the TBD has only slightly shifted towards the Lon domain (**Fig. 2c**).

**Table 1.** Conformational state distribution of CRBN following compound treatment, order of addition, and incubation time.

| Order of treatment | Incubation time (minutes) |  | Open % | Intermediate % | Closed % | Frustrated % |
| --- | --- | --- | --- | --- | --- | --- |
|  | with inhibitor | with IMiDs |  |  |  |  |
| VVD-105917 alone | 2 |  | 100 | - | - | - |
| Iberdomide alone (Watson et al., 2022) |  | 2 | 50 | - | 50 | - |
| Mezigdomide alone (Watson et al., 2022) |  | 2 | - | - | 100 | - |
| VVD-105917 + Iberdomide | 2 | 2 | 100 | - | - | - |
| VVD-105917 + Iberdomide | 10 | 30 | 100 | - | - | - |
| Iberdomide + VVD-105917 | 30 | 10 | 52 | 38.5 | 9.5 | - |
| VVD-105917 + Mezigdomide | 10 | 30 | 100 | - | - | - |
| Mezigdomide + VVD-105917 | 30 | 10 | 18 | 15 | 23 | 44 |
| Mezigdomide + VVD-105917 | 60 | 10 | 12 | 9 | 14 | 65 |
| Mezigdomide + VVD-105917 | 360 | 60 | 12 | 50 | 15 | 23 |

Notably, in the intermediate state, we observe density corresponding to bound VVD-105917; however, it is slightly shifted relative to its position in the fully open conformation, consistent with partial domain rearrangement (**Fig. 2d**). The model built from this open-conformation population revealed density for both compounds bound simultaneously, although density for iberdomide itself was resolved only for the glutarimide-isoindolinone core engaging the tri-Trp pocket; the appended aryl-morpholine substituent was not ordered, consistent with the absence of a neosubstrate to stabilize this portion of the molecule.

We observed a notable difference in the sensor loop compared to the previously determined IMiD-bound open-state conformer of cereblon (PDB ID: 8cvp) ^9^. Whereas this loop was thought to be largely flexible in the open IMiD-bound conformer and organizing as a β-hairpin upon CRBN closure, our dual-liganded open-state map contained density sufficient to model much of this previously missing segment (residues 340-352) (**Fig. 2b**). This density is consistent with a previously proposed interaction between the sensor loop and DDB1^9^ and suggests that stabilization of the long-lived open conformation by VVD-105917 enables visualization of this interaction with greater clarity. While the loop remains conformationally heterogeneous, these data suggest that it is not completely disordered in the open conformation.

### VVD-105917 counteracts CRBN closure even under a potent CELMoD

Prior work has shown that IMiDs/CELMoDs bind the orthosteric pocket of CRBN in either open or closed conformations and that the shift in the population distribution along the open-closed equilibrium is influenced by chemical structure, rather than reversible binding ^9^. A key unresolved question has been whether the liganded closed state is effectively “locked,” or whether it can reversibly reopen on experimentally relevant timescales. Since the Cys287 allosteric pocket is occluded in the fully closed state, the results of our sequential-incubation cryo-EM experiments suggest that liganded closed CRBN will revisit the open conformation.

To further assess how a more potent driver of CRBN closure behaves in the presence of VVD-105917, we repeated the sequential-incubation experiment using mezigdomide, a CELMoD shown to drive CRBN to full closure ^9^. As previously observed with iberdomide, adding mezigdomide to VVD-105917-bound CRBN did not promote CRBN closure (**Table 1**), showcasing the VVD compound’s potent inhibition of the system. We next added VVD-105917 to fully close mezigdomide-bound CRBN. Despite this strong closure bias, addition of VVD-105917 to mezigdomide-bound CRBN for 30-60 minutes dramatically redistributed the conformational ensemble, with only 15-25% of the particles remaining in the closed state. Of the remaining particles, 12-18% were observed in the open conformation; 10-15% occupying an intermediate state; and 40-60% populating a distinct “frustrated” intermediate (**Table 1**). The frustrated class adopts a near-closed architecture in which the VVD-105917 pocket is occluded. Extending the incubation (1 h mezigdomide followed by 6 h VVD-105917) did not drive complete reopening, but instead progressively enriched intermediate states, suggesting that under these conditions the system approaches a kinetic steady state among open, intermediate, and (near-) closed conformations rather than a fully reopened end-point. These structural data directly correlate with functional outcomes: treatment with 10 µM VVD-105917 effectively abolishes iberdomide-mediated IKZF3 degradation (**Fig. 1e**), demonstrating that the allosteric inhibitor-stabilized open state is incompatible with IMiD activity.

## Discussion

The discovery that Cys287 resides within an allosteric pocket capable of locking CRBN in its open conformation has notable mechanistic implications. The recent study by Dippon et al. ^13^ identified a structurally distinct ligand that binds this same allosteric region yet can produce the opposite functional outcome: enhancement of IMiD-induced neosubstrate degradation. Together, these results point to a sign-switchable allosteric site: the same allosteric pocket on CRBN can behave as either an activator or an inhibitor depending on the chemical structure and binding mode of the occupying ligand, a phenomenon that is shared with several recent disclosures of allosteric pockets ^17–19^. Our work further demonstrates that covalent engagement at Cys287 can override orthosteric IMiD binding, enforcing an open, inactive state regardless of the presence of closure-promoting ligands.

These findings indicate that ligand-induced regulation at the Cys287 allosteric site is highly tunable: depending on the chemistry and binding mode, occupancy of the same pocket can either potentiate or suppress IMiD/CELMoD-driven neo-substrate degradation. Activating ligands likely promote cooperative stabilization of the closed CRBN state or enhance the efficiency of neomorphic interface formation, thereby amplifying substrate recruitment. In contrast, inhibitory ligands such as VVD-105917 bias CRBN toward the open conformation, preventing formation of the IMiD-dependent binding surface required for degron recognition. One can thus speculate that covalent C287 ligands that potentiate rather than inhibit neosubstrate degradation may someday be identified for therapeutic pursuits.

The VVD-105917-bound open structure revealed two further layers of conformational nuance. A previously unresolved N-terminal segment positioned between the Lon domain and DDB1 (**Fig. 2a, Supplementary Fig. 2**) suggests another allosteric interface, though whether this plays a functional role or simply reflects open-state stabilization remains to be determined. Separately, within open-conformation particles from the iberdomide-then-VVD-105917 dataset, the sensor loop showed partial ordering despite the domain-level conformation remaining open; rather than an independent event, this likely represents an early step along the same pathway toward the intermediate and closed states, with iberdomide able to locally reorganize the sensor loop even while VVD-105917 keeps the domain open -a state not accessible with either ligand alone. Together, these findings add further layers to the tunable allostery described above. Collectively, these findings reveal an unexpected degree of conformational plasticity within CRBN and establish cryptic allosteric sites as a versatile design space for both enhancing and suppressing targeted protein degradation.

## Material & Methods

### Chemoproteomic screening and target engagement measurements

Jurkat, MM.1S, or PBMC cell pellets were lysed and analyzed for cysteine reactivity as previously described ^16^. Briefly, cell pellets were lysed by probe sonication in DPBS. Protein concentration was normalizd to 1mg/mL and lysates were treated with compound for 1 h at room temperature prior to addition of IA-dTB probe (custom synthesis, WuXi App Tec) for 1 h. For some experiments, lysates were denatured via addition of 9M urea prior to probe-labeling. Samples were treated with benzonase nuclease (Millipore Sigma, catalogue no. 70746-3) and then subjected to acetone precipitation. Precipitated protein was resuspended in 9 M urea, 50 mM ammonium bicarbonate, 10 mM dithiothreitol, alkylated with IAA, and desalted with Zeba desalting plates (Thermo Fisher Scientific, catalogue no. 89807). Following trypsin digestion, cysteine-containing peptides were captured using 5% high-capacity streptavidin agarose. Eluted peptides were dried and resuspended in 3% acetonitrile and 0.1% formic acid for mass spectrometry analysis. Engagement (at a given concentration) was calculated by comparing compound-treated samples to DMSO-treated controls.

### Liquid chromatography–tandem mass spectrometry analysis

Probe-labelled peptides were analyzed via LC-MS/MS essentially as previously described ^16^. Targeted measurements were performed via parallel reaction monitoring (PRM) mass spectrometry using product ion scan mode on a Thermo Exploris orbitrap mass spectrometer and data were analyzed using Skyline V21.2.0.369 (MacCoss Lab, University of Washington). Global cysteine profiling of VVD-105917 was performed essentially as previously described ^18^. Briefly, after isolating probe-labeled peptides, samples were labeled by addition of TMTpro 16plex reagent (Thermo Fisher Scientific, A44520) and quenched by the addition of 6.5uL of 5% hydroxylamine. Samples were then combined, desalted, and the resulting peptides separated into 12 fractions via high-pH reversed-phase chromatography and subjected to MS analysis on a Lumos Orbitrap mass spectrometer. Data were analyzed with the MassPIKE software package (GFY Core 101 Version 3.4). Searched data were further filtered using in-house scripts as previously described.

### Detection of protein degradation via nanoluciferase and immunoblot

Targeted degradation assays were performed in HEK293LX cells plated at 750,000 cells per well in 6-well plates. When cells reached approximately 40% confluency they were transfected using Fugene HD (Promega E2311) and Lonza-GFP (V1435; 3 µg) as a carrier alongside either BRD4-NanoLuc (V1546) or FKBP12^F36V^-NanoLuc (V1330; 1.5 ng) for a total of 3 µg DNA per well. Twenty-four hours post-transfection, cells were dissociated with accutase and reseeded at approximately 15,000 cells per well in 384-well plates in 20 µL volume of culture medium, then allowed to adhere for 2–3 hours at 37°C, 5% CO_2_. Cells were then pre-treated with a 10-point dose-response of VVD-105917 for 2 hours. Following pre-treatment, targeted degradation was initiated by addition of either 2 µM dBET1 or 10 nM dTAG-13 with DMSO serving as vehicle control. After 4 hours, 24 µL of HiBiT lytic substrate (Promega, N3030) was added to each well and luminescence was measured after 10 minutes. Data are expressed as percentage of DMSO control.

### Synthesis of VVD-105917

A mixture of 3-bromo-5-hydroxybenzonitrile (1 g, 5.05 mmol), (2-117 methoxyphenyl)boronic acid (0.767 g, 5.0 mmol), potassium carbonate (0.698 g, 5.0 mmol), and Pd(dppf)Cl2 (185 mg) in acetonitrile (4 mL), 1,4-dioxane (4 mL), and water (2 mL) was degassed and purged with N2 three times, then stirred at 100°C for 6 h under N2. The mixture was partitioned between water (15 mL) and EtOAc (3 120 × 10 mL); the organic phase was dried over sodium sulfate, filtered, concentrated, and purified by column chromatography (silica, petroleum ether/ethyl acetate, 10:1 to 1:1) to give 3-hydroxy-5-(2-122 methoxyphenyl)benzonitrile (1.0 g, 88% yield). To a solution of 3-hydroxy-5-(2-methoxyphenyl)benzonitrile (0.62 g, 2.75 mmol), tert-butyl 3-fluoro-4-hydroxypiperidine-1-carboxylate (0.604 g, 2.75 mmol), and PPh3 (1 g, 5.51 mmol) in THF (31 mL) at 15°C under nitrogen was added DIAD (834 mg, 5.51 mmol); the mixture was stirred at 60°C for 16 h, then partitioned between water (40 mL) and EtOAc (2 × 30 mL). The organic phase was washed with brine, dried over sodium sulfate, filtered, concentrated, and purified by preparative TLC (petroleum ether/EtOAc, 3:1) to give tert-butyl 4-[3-cyano-5-(2-methoxyphenyl)phenoxy]-3-fluoropiperidine-1-carboxylate (0.50 g, 43% yield) as a yellow oil. Treatment with HCl in EtOAc (10 mL, 4 M) in dichloromethane (1 mL) at room temperature for 30 min, followed by concentration under reduced pressure, gave 3-[(3-fluoropiperidin-4-yl)oxy]-5-(2-methoxyphenyl)benzonitrile, which was carried forward without further purification. To a solution of this amine (340 mg, 1.0 mmol) in dichloromethane (19 mL) was added triethylamine (0.44 mL, 3.1 mmol); the mixture was stirred at 0°C for 30 min, then treated with acryloyl chloride (104 mg, 1.1 mmol) and stirred at room temperature for 16 h. The mixture was treated with aqueous ammonium chloride (4 mL) and ethyl acetate (2×4 mL); the organic phase was washed with brine, dried over sodium sulfate, and concentrated, and the residue was purified by supercritical fluid chromatography (SFC) to separate the diastereomers and provide VVD-105917, 5-(((3S,4R)-1-acryloyl-3-fluoropiperidin-4-yl)oxy)-2’-methoxy-[1,1’-biphenyl]-3-carbonitrile. ^1^H NMR (400 MHz, CDCl3) d 1.91 (br s, 1H), 2.18 (br s, 1H), 3.45–3.83 (m, 2H), 3.84 (s, 3H), 3.89–4.10 (m, 2H), 138 4.57–4.80 (m, 2H), 5.75 (dd, J = 10.5, 1.8 Hz, 1H), 6.32 (br d, J = 16.3 Hz, 1H), 6.59 (dd, J = 16.8, 10.5 Hz, 1H), 6.99–7.09 (m, 2H), 7.17 (dd, J = 2.5, 1.3 Hz, 1H), 7.27–7.30 (m, 1H), 7.32–7.43 (m, 2H), 7.49 (t, J = 1.3 Hz, 1H).

### Protein expression and purification

#### DDB1^ΔBPB^/CRBN expression and purification

The DDB1^ΔBPB^/CRBN complex was expressed and purified as previously described ^9^. Briefly, 6XHis-tagged HsCRBN (residues 1–442) and HsDDB1^ΔBPB^, (396-705 truncation with addition of a GNGNSG linker between BPA and BPC) were cloned in the Bac-to-Bac Baculovirus Expression System. Bacmids were generated in DH10 cells following the manufacturer’s instructions (ThermoFisher), and 1 µg of each vector was co-transfected into Spodoptera frugiperda (Sf9) cells at a density of 1.5 *×* 10^6^ cells/ml to generate recombinant baculovirus. Viral stocks were amplified to the second generation (P2), and fresh Sf9 cells (2 *×* 10^6^ cells/ml) were co-infected with 3% (v/v) of each P2 virus in the presence of 50 µM ZnSO_4_. Infected cells were harvested 48 h post-infection, flash-frozen in liquid nitrogen, and stored at –80 °C. The cell pellet was resuspended in buffer containing 50 mM Tris-HCl (pH 7.5), 300 mM NaCl, 10 mM imidazole, 10% glycerol, 3 mM TCEP, 1X Protease Inhibitor Cocktail (Roche), and 40,000 U Benzonase (Novagen), and sonicated for 30 s. Lysate was clarified by high-speed centrifugation at 108,800 g for 30 min, and clarified lysate was incubated with TaLon^R^ metal affinity resin (Takara) for 1 h. Complex was eluted with buffer containing 250 mM imidazole, and diluted to 150 mM NaCl for further purification over a MonoQ ion exchange column (GE Healthcare). The MonoQ column was washed with ten column volumes of 50 mM Tris-HCl (pH 7.5), 150 mM NaCl, 3 mM TCEP, followed by equilibration with ten column volumes of 50 mM Bis-Tris (pH 6.0), 150 mM NaCl, 3 mM TCEP, sample was loaded and subjected to elution with a NaCl gradient 0.15M-1M using the Bis-Tris buffer. The DDB1^ΔBPB^/CRBN peak was eluted at 200 mM NaCl. This peak was collected and further purified by size-exclusion buffer containing 10 mM HEPES pH 7.5, 240 mM NaCl, and 3 mM TCEP. Samples were concentrated, aliquoted, flash-frozen, and stored at -80 °C.

#### Mutant IKZF1 (ZF2-3) protein purification

Mutant 140-196^Q146A, G151N^) Ikaros protein zinc fingers (ZF2-3) were cloned into pET28A expression vectors and purified as 6XHis-MBP (maltose binding protein) tag proteins from *E. coli* Rosetta cells (Novagen). Cells were induced with 0.5 mM isopropyl β-D-1-thiogalactopyranoside (IPTG) at OD^600^ 0.6 for 18 h at 16 °C. Cells were pelleted at 4000g, flash frozen with LN_2_, and stored at -80°C. Cells were thawed and resuspended in buffer containing 50 mM Tris-HCl (pH 7.5), 200 mM NaCl, 10 mM imidazole, 10% glycerol, 3 mM TCEP, 1X Protease Inhibitor Cocktail (Roche), 25000U/ml DNAseA, 1 mM PMSF, and 50 µM ZnCl2 and subjected to metal affinity purification using TaLonR resin as described above, up to the binding step. TEV protease was added at a 1:100 ratio to the bead-bound proteins and incubated overnight at 4°C to cleave the His_6_-MBP tag. The flow-through, devoid of both the tag and TEV (which was also 6XHis-tagged), was collected, concentrated, and further purified by size-exclusion buffer containing 10 mM HEPES pH 7.5, 240 mM NaCl, 3 mM TCEP, and 50 µM ZnCl2. The final sample was concentrated, aliquoted, flash-frozen, and stored at -80 °C.

### Sample preparation for cryo-EM

DDB1^ΔBPB^/CRBN(40 µM) was incubated with VVD-105917 (80 µM) in dilution buffer containing 10 mM HEPES, pH 7.0, 240 mM NaCl, and 3 mM TCEP for 2 min at 4 °C. Final DMSO and glycerol concentrations were maintained below 1% (v/v). Quantifoil 300-mesh R1.2/1.3 UltrAuFoil holey gold grids were plasma cleaned for 15 s in a Solarus plasma cleaner (Gatan) using a 75% argon and 25% oxygen atmosphere at 15 W under ambient vacuum. To limit complex dissociation, each grid was preincubated with 4 µl of 10 µM CRBN-agnostic mutant Ikaros residues 140-196^Q146A, G151N^ for 1 min and back-blotted with Whatman No. 1 filter paper. Immediately before plunge-freezing, 1 µl of complex was diluted tenfold into 5 µl of dilution buffer supplemented with 0.002% lauryl maltose neopentyl glycol and 4 µl of mutant Ikaros solution. A 4-µl aliquot of this mixture was applied to the grid and blot-plunged using a manual plunge freezer in a 4 °C cold room at greater than 95% relative humidity. Grids were blotted for approximately 3 s after the sample droplet stopped expanding and immediately plunged into liquid ethane cooled by liquid nitrogen. For samples containing DDB1^ΔBPB^/CRBN, VVD-105917, and an IMiD, either VVD-105917 or the IMiD was added first and incubated for the specified interval before addition of the second compound; samples were then incubated for the second specified interval before freezing, as detailed in (**Table 1**). DDB1^ΔBPB^/CRBN, VVD-105917, and the IMiD were combined at a molar ratio of 1:2:2.5, respectively.

### Cryo-EM data acquisition

Cryo-EM data were collected on a Thermo-Fisher Talos Arctica transmission electron microscope operating at 200 keV using parallel illumination conditions ^20,22^. Micrographs were acquired using a ThermoFischer Scientific Falcon 4i direct electron detector, operated in electron counting mode, applying a total electron exposure of 50 e-/Å 2. The Thermo Fisher Scientific EPU version 3.9.1.8 data collection software was used to collect micrographs at 190,000x nominal magnification (0.731 Å /pixel at the specimen level) with a nominal defocus set to 0.8 µm -1.8 µm under focus. Stage movement was used to target the center of either 4 or 16 1.2 µm holes for focusing, and image shift to 8 µm was utilized to acquire high-magnification images. The cryoSPARC Live software package was used for real-time image analysis and assessment. Patch motion correction and patch CTF estimation with default parameters were used during collection to monitor data quality. Only images with CTF resolution estimates of 2 to 4 Å were selected for further processing. The cryoSPARC Blob picker was used to select particles with a diameter set to 110-140 Å, and were extracted with a 300-pixel extraction box size, downscaled to 100 pixels for analysis. Detailed dataset specifics are available in **Supplementary Table 1**. In brief, 4,553 micrographs were collected for DDB1^ΔBPB^/CRBN + VVD-105917, 9,019 micrographs for DDB1^ΔBPB^/CRBN + VVD-105917 + iberdomide (2 minutes incubation with each drug), 2,498 micrographs for DDB1^ΔBPB^/CRBN + VVD-105917 + iberdomide (10 minutes incubation with VVD-105917 and 30 minutes incubation with iberdomide), 3,002 micrographs for DDB1^ΔBPB^/CRBN + iberdomide + VVD-105917 (10 minutes incubation with iberdomide and 30 minutes incubation with VVD-105917), 2,492 micrographs for DDB1^ΔBPB^/CRBN + VVD-105917 + mezigdomide (10 minutes incubation with VVD-105917 and 30 minutes incubation with mezigdomide), 3,349 micrographs for DDB1^ΔBPB^/CRBN + mezigdomide + VVD-105917 (10 minutes incubation with mezigdomide and 30 minutes incubation with VVD-105917), 2,859 micrographs for DDB1^ΔBPB^/CRBN + mezigdomide + VVD-105917 (10 minutes incubation with mezigdomide and 60 minutes incubation with VVD-105917), and 2,832 micrographs for DDB1^ΔBPB^/CRBN + mezigdomide + VVD-105917 (60 minutes incubation with mezigdomide and 360 minutes incubation with VVD-105917).

### Cryo-EM data processing and image analysis

A representative image-processing workflow for all datasets is shown in (**Supplementary Fig. 3**). Processing was performed in cryoSPARC. At the end of each acquisition session, accepted exposures and particles from cryoSPARC Live were exported to a cryoSPARC workspace. Accepted exposures were subjected to blob picking using the parameters described above, and the resulting particles underwent 2D classification for initial cleanup. Selected particles from the best 2D classes were used to generate three ab initio maps representing open, closed, and junk classes. These maps served as references for heterogeneous refinement of all particles using default parameters and C1 symmetry. For subsequent datasets, particles exported from cryoSPARC Live were classified directly by heterogeneous refinement against these conformational references. Particles assigned to open or closed states were then subjected to 3D classification without alignment using PCA initialization, an O-EM initial learning rate of 0.1, an O-EM half-life of 100 iterations, a class-similarity value of 0.1, a 6-Å filter resolution, five O-EM epochs, and forced hard classification. One or two classes derived from the closed-state particles were assigned as intermediate states. Particle subsets for final reconstruction were selected based on homogeneity across the full particle, map-feature quality, and estimated resolution at the FSC 0.143 criterion, then re-extracted in the full 300-pixel box. These particles were refined using non-uniform refinement with one extra final pose, a 6-Å initial low-pass resolution, per-particle defocus refinement, and per-group CTF parameter refinement. Reference-based motion correction, alignment improvement, and additional non-uniform refinement were performed to obtain the final maps. The CRBN region was locally refined using a soft mask shown in **Supplementary Fig. 3**, with a 3° standard-deviation prior on rotation, a 2-Å standard-deviation prior on shift, a 15° rotation search, a 10-Å shift search, recentering of rotations and shifts after each iteration, two extra final poses, and a 6-Å initial low-pass resolution. Local-resolution maps were calculated in cryoSPARC and are shown in (**Supplementary Figs. 4 and 5**).

### Atomic model building and refinement

Model building and refinement began from the the published DDB1^ΔBPB^/CRBN cryo-EM structure (PDB 8CVP). Coordinates and restraints for VVD-105917 were generated using the PDB Chemical Sketch Tool ^29^, and those for iberdomide were generated using eLBOW in PHENIX ^23^. Newly resolved density in both structures from this study (PDB 38MG and 38MJ) was interpreted de novo in Coot with reference to the CRBN sequence. For 38MG, the register of the N-terminal segment was anchored by distinctive side-chain density for Asp63, Met64, Glu66, Phe67, and Arg70. CRBN residues 71–77 were not modeled because the corresponding density lacked sequence-specific features and was not clearly resolved at a high map threshold. In the VVD-105917–iberdomidebound structure 38MJ, previously unresolved density for the sensor loop, which is disordered over residues 340–356 in PDB 8CVP, allowed residues 340–352 to be built de novo, with a short internal break at residue 347; residues 353–356 remained unresolved and were not modeled. Iberdomide density was resolved only for the glutarimide–isoindolinone core engaging the tri-Trp pocket; the unresolved aryl-morpholine substituent was therefore omitted. Iterative model building and refinement were performed using PHENIX v2.0rc1-5617^24^ and Coot 0.9.8.95 EL ^25^ until agreement between each model and its map was achieved. The first cycle of PHENIX real-space refinement included global minimization, rigid-body refinement, local grid search, atomic displacement-parameter refinement, and morphing. Refinement was run for 100 iterations and five macrocycles with target bond and angle root-mean-square deviations of 0.01 Å and 1.0°, respectively, with secondary-structure and Ramachandran restraints enabled. In some cases, models were improved using maps generated with EMReady2^26^ or composite maps combining non-uniform and local refinements. Validation was performed against combined, unsharpened non-uniform and local-refinement maps, with statistics reported in (**Supplementary Table 2**). UCSF Chimera 1.7.1^27^ and ChimeraX Daily ^28^ were used to inspect the reconstructions and atomic models and to prepare figures. **Supplementary Tables 1 and 2** report data-collection, refinement, and validation statistics.

## Data availability

CryoEM maps and associated atomic model were deposited to the Electron Microscopy Databank (EMDB) and the Protein Databank (PDB), respectively, with the following EMDB and PDB IDs: DDB1^ΔBPB^/CRBN + VVD-105917 (2 minutes incubation) complex: CRBN^open^ - EMD-78925 (composite map), EMD-78923 (consensus map), EMD-78924 (focus map), PDB ID 38MG. DDB1^ΔBPB^/CRBN + VVD-105917 (2 minutes incubation) + Iberdomide (2 minutes incubation) complex: CRBN^open^ - EMD-78926 (consensus map), EMD-78927 (focus map). DDB1^ΔBPB^/CRBN + VVD-105917 (10 minutes incubation) + Iberdomide (30 minutes incubation) complex: CRBN^open^ - EMD-78928 (consensus map), EMD-78929 (focus map), EMD-78932 (composite map), and PDB ID 38MJ. DDB1^ΔBPB^/CRBN + Iberdomide (10 minutes incubation) + VVD-105917 (30 minutes incubation) complex: CRBN^open^ - EMD-78930 (consensus map), EMD-78931 (focus map), CRBN^intermediate^ - EMD-78933 (consensus map), EMD-78934 (focus map), and CRBN^closed^ - EMD-78966 (consensus map), EMD-78967 (focus map). DDB1^ΔBPB^/CRBN + VVD-105917 (10 minutes incubation) + Mezigdomide (30 minutes incubation) complex: CRBN^open^ - EMD-78969 (consensus map), EMD-78970 (focus map). DDB1^ΔBPB^/CRBN + Mezigdomide (10 minutes incubation) + VVD-105917 (30 minutes incubation) complex: CRBN^open^ - EMD-78971 (consensus map), EMD-78972 (focus map), CRBN^intermediate^ - EMD-78973 (consensus map), EMD-78974 (focus map), CRBN^closed^ - EMD-78975 (consensus map), EMD-78976 (focus map), and CRBN^frustrated^ - EMD-78977 (consensus map), EMD-78978 (focus map). DDB1^ΔBPB^/CRBN + Mezigdomide (10 minutes incubation) + VVD-105917 (60 minutes incubation) complex: CRBN^open^ - EMD-78986 (consensus map), EMD-78987 (focus map), CRBN^intermediate^ - EMD-78982 (consensus map), EMD-78983 (focus map), CRBN^closed^ - EMD-78988 (consensus map), EMD-78989 (focus map), CRBN^frustrated1^ - EMD-78990 (consensus map), EMD-78997 (focus map), and CRBN^frustrated2^ - EMD-78998 (consensus map), EMD-78999 (focus map). DDB1^ΔBPB^/CRBN + Mezigdomide (60 minutes incubation) + VVD-105917 (360 minutes incubation) complex: CRBN^open^ - EMD-79011 (consensus map), EMD-79012 (focus map), CRBN^intermediate^ - EMD-79002 (consensus map), EMD-79004 (focus map), CRBN^closed^ - EMD-79005 (con- sensus map), EMD-79008 (focus map), and CRBN^frustrated^ - EMD-79009 (consensus map), EMD-79010 (focus map).

## Acknowledgments

We thank Chris Zerio, Steffen Bernard, and Jan Hannes Schäfer for helpful discussions in model interpretations. We thank JC Ducom at Scripps Research High Performance Computing and Charles Bowman at Scripps Research for computational support, as well as Will Lessin at the Scripps Research Electron Microscopy Facility for microscopy support. GCL is supported by NIH grant GM143805, and the work used equipment supported by NIH grant S10OD032467. ZR is supported with American Cancer Society postdoctoral fellowship.

## Author contributions

G.C.L., G.M.S., and M.P.P. supervised the project. G.C.L. obtained funding for the structural studies. Z.R. conceived, designed, and performed the cryo-EM studies and analyzed the structural data. B.E.N., K.T.S., and K.A.B. performed proteomic and degradation studies. D.A.W. designed VVD-105917 and oversaw its synthesis and characterization Z.R. drafted the manuscript with input from all authors. All authors reviewed and edited the manuscript.

## Competing interests

B.E.N., M.P.P., and G.M.S. are current employees of Vividion Therapeutics. K.A.B. was employee of Vividion Therapeutics when the research was conducted. G.C.L. is a current employee of Bristol Myers Squibb. The remaining authors declare no competing interests.

## Supporting Information

**Supplementary Figure 1.**
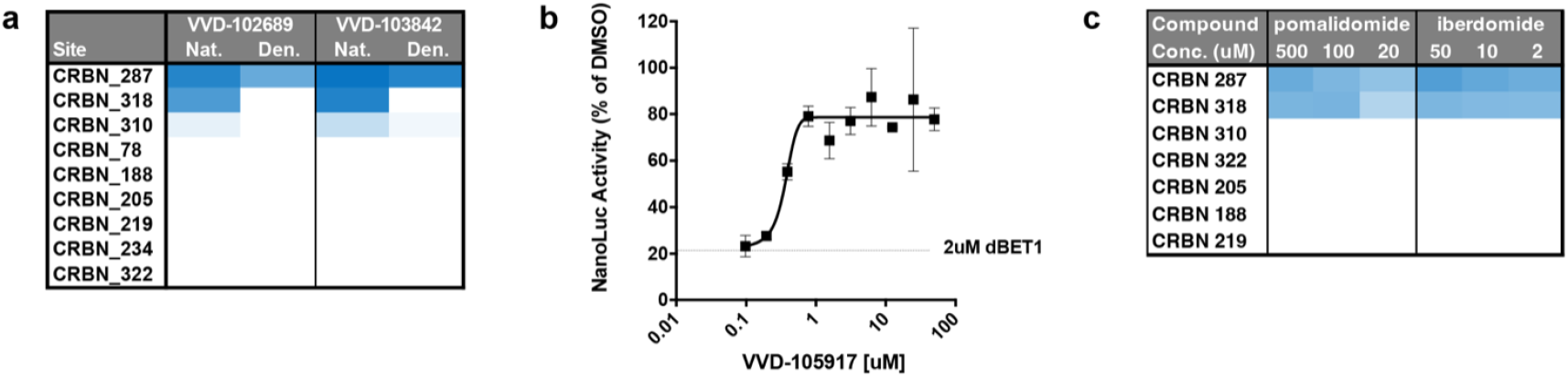
Chemoproteomic characterization of CRBN cysteine engagement and IMiD competition. **a**, Probe labeling under native and denaturing conditions reveals covalent engagement of Cys287. PBMC lysate was treated with VVD-102689 or VVD-103842, followed by IA-dTB probe labeling under native or denaturing conditions. The heat map shows inhibition of probe labeling for each CRBN cysteine-containing peptide; darker blue indicates greater probe competition. Inhibition at Cys287 is preserved under denaturing conditions, supporting direct covalent modification, whereas competition at Cys318 is lost upon denaturation. **b**, VVD-105917 inhibits CRBN-dependent degradation of BRD4 mediated by dBET1. HEK293LX cells expressing BRD4-NanoLuc were pretreated with VVD-105917 for 2 h before addition of 2 µM dBET1. BRD4 protein levels were assessed after 4 h by HiBiT luminescence. **c**,The IMiDs pomalidomide and iberdomide reduce IA-dTB probe labeling at Cys287 and Cys318. CRBN-expressing PBMC lysates were treated with pomalidomide (500, 100, or 20 µM) or iberdomide (50, 10, or 2 µM) before probe labeling under native conditions. The heat map shows inhibition of probe labeling at each CRBN cysteine; darker blue indicates greater probe competition. Both IMiDs partially suppress labeling at Cys287 and Cys318, suggesting that the IMiD-bound closed conformation of CRBN restricts access to the Cys287 allosteric pocket.

**Supplementary Figure 2.**
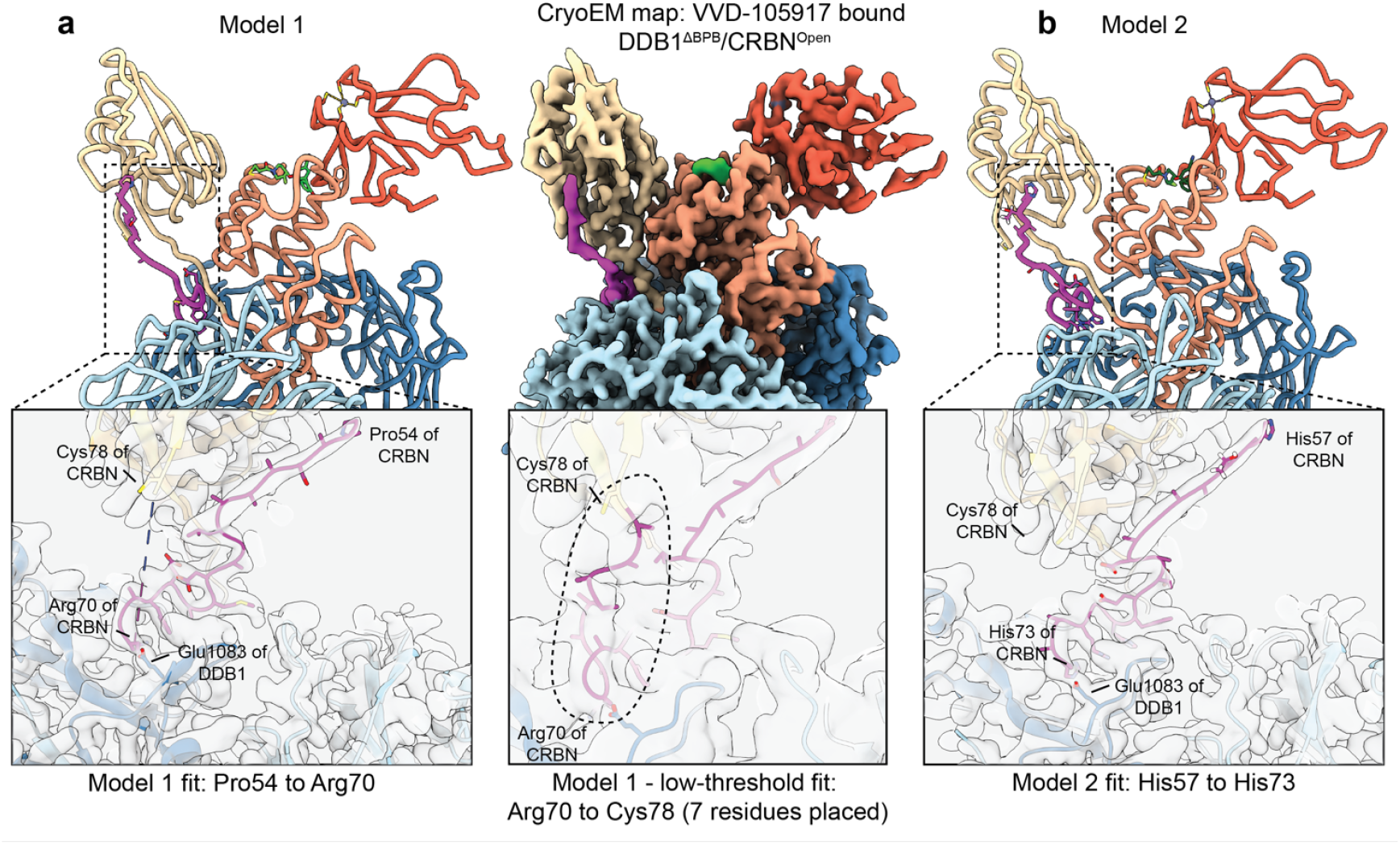
Two candidate sequence registers for the newly identified CRBN N-terminal density. In the reference open-conformation structure (PDB 8CVP) and other cryo-EM models (PDB 8D7X and 8D7Y), the ordered CRBN chain begins at Met64; available crystal structures (PDB 6H0F, 6BNB, and 7U8F) begin at Gly69. Thus, the preceding 63 residues have not been characterized in a published open-conformation structure. The density in the VVD-105917-bound map is shown in the center panel (composite cryo-EM map of the DDB1^ΔBPB^/CRBN complex, colored as in Fig. 2, with the uncharacterized density in magenta) and supports two alternative sequence registers. **a**, Model 1. The N terminus is modeled from Pro54 (residues 54–70), extending downward into the new density. Distinctive side-chain densities for Asp63, Met64, Glu66, Phe67, and Arg70 anchor the register, and Arg70 is positioned to interact with DDB1 Glu1083. **b**, Model 2. The N terminus is modeled from His57, extending downward into the same density. Distinctive side-chain densities for Asp63, Glu66, Phe67, Arg70, and His73 anchor the register, and His73 is positioned to interact with DDB1 Glu1083. In both models, residues beyond the last modeled position are unresolved at the standard map threshold; weak density at a lower contour level is consistent with the chain looping upward to rejoin the established chain at Cys78. The gap length favors Model 1, which requires seven unmodeled residues (71–77) to connect to Cys78, whereas Model 2 requires four (74–77).

**Supplementary Figure 3.**
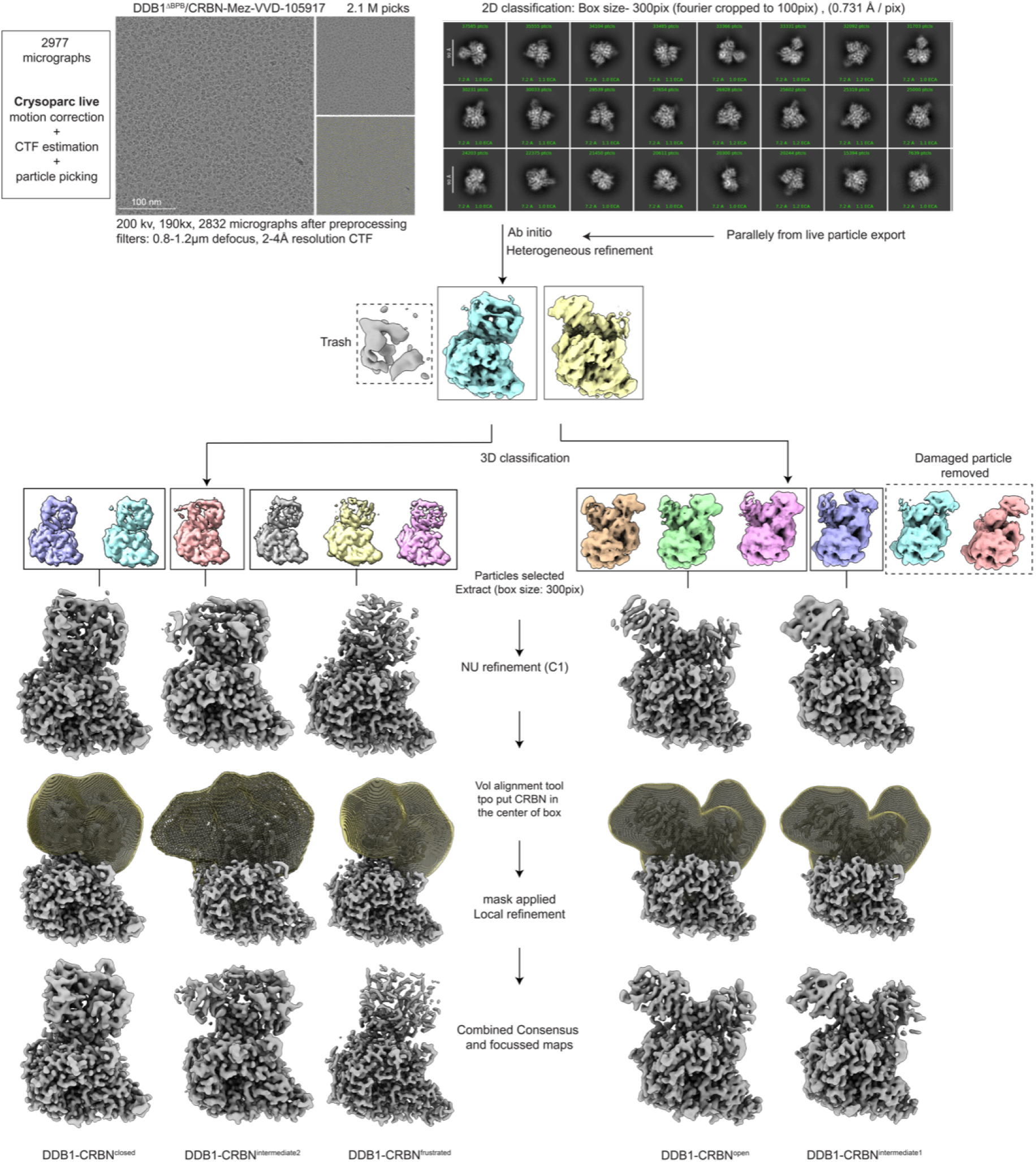
CryoEM image analysis workflow. Representative processing workflow described for DDB1^ΔBPB^/CRBN treated with mezigdomide and VVD-105917. For all datasets, motion correction, CTF estimation, particle picking, and extraction were performed in real time using cryoSPARC Live. The data were then imported into cryoSPARC for further processing, including 2D classification, ab initio reconstruction, heterogeneous refinement, and 3D classification, as detailed in the methods section. Ab initio 3D models were generated from the particles contributing to the best class averages and used to perform 3D classification to identify conformational and compositional heterogeneity. Particles belonging to similar classes were combined and subjected to non-uniform refinement. Local regions of CRBN were refined using local refinement jobs. Final reconstructions for five conformational states are shown at the bottom.

**Supplementary Figure 4.**
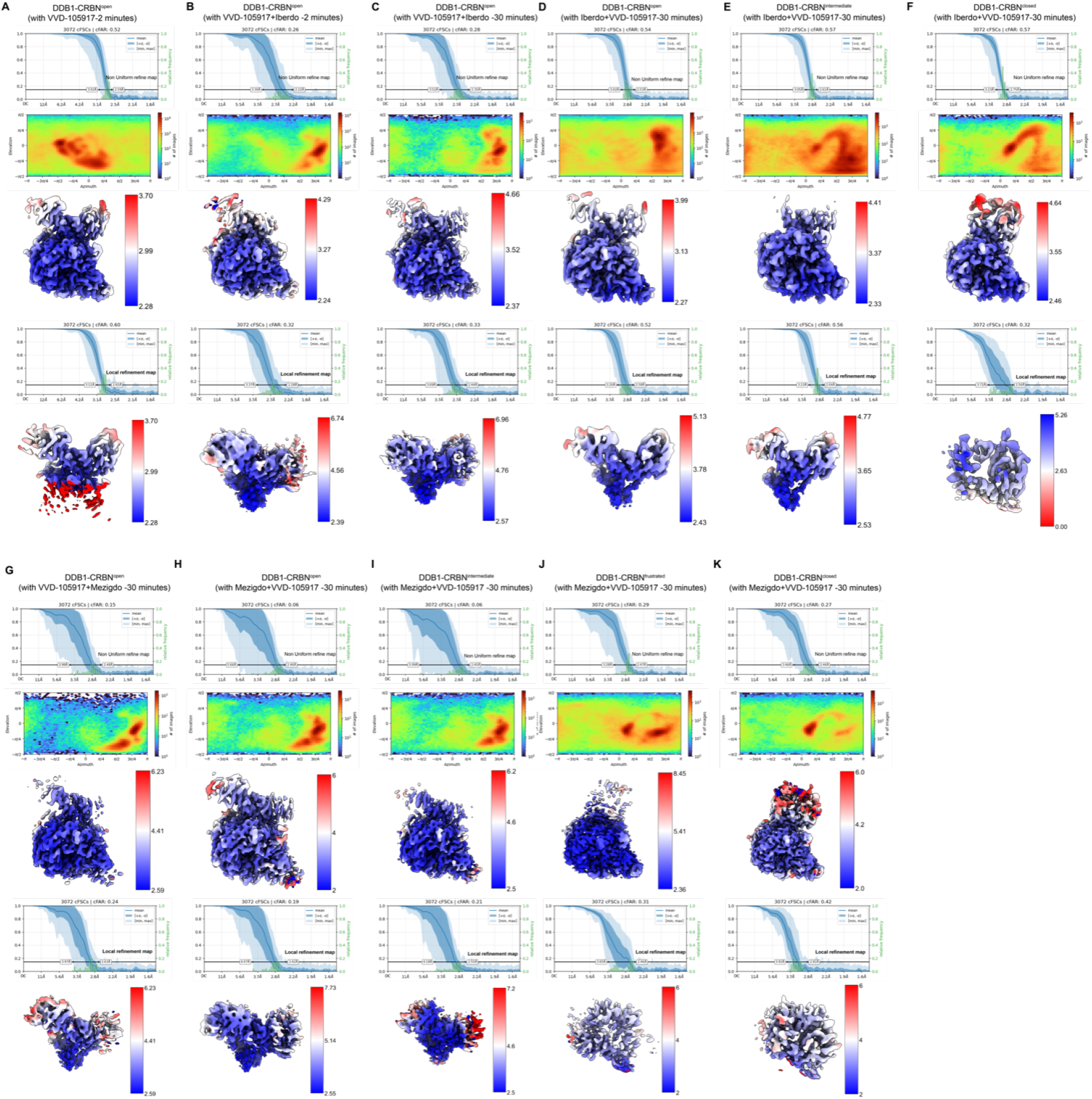
Resolution, angular distribution, and local resolution assessment of cryo-EM reconstructions. **(a-k)** Cryo-EM data quality metrics for each reconstruction. Each panel contains, from top to bottom: conical Fourier shell correlation (cFSC) curve (blue plot), with overlaid histogram of directional resolutions (green); the angular distribution plot; full reconstruction colored by local resolution; cFSC curve for the masked local refinement of the CRBN domain; and locally refined CRBN domain colored by local resolution. The reconstructions shown are: **a**, DDB1^ΔBPB^/CRBN^open^ after 2-minute incubation with VVD-105917. **b**, DDB1^ΔBPB^/CRBN^open^ after sequential 2-minute incubation with VVD-105917 followed by iberdomide. **c**, DDB1^ΔBPB^/CRBN^open^ after 10-minute incubation with VVD-105917 followed by 30-minute incubation with iberdomide. **d-f**, DDB1^ΔBPB^/CRBN^open^ **d**, DDB1^ΔBPB^/CRBN^intermediate^ **e**, and DDB1^ΔBPB^/CRBN^closed^ **f**, after 10-minute incubation with iberdomide followed by 30-minute incubation with VVD-105917. **g**, DDB1^ΔBPB^/CRBN^open^ after 10-minute incubation with VVD-105917, followed by 30-minute incubation with mezigdomide. **h-k**, DDB1^ΔBPB^/CRBN^open^ **h**, DDB1^ΔBPB^/CRBN^intermediate^ **i**, DDB1^ΔBPB^/CRBN^frustrated^ **j**, and DDB1^ΔBPB^/CRBN^closed^ **k**, after 10-minute incubation with mezigdomide followed by 30-minute incubation with VVD-105917.

**Supplementary Figure 5.**
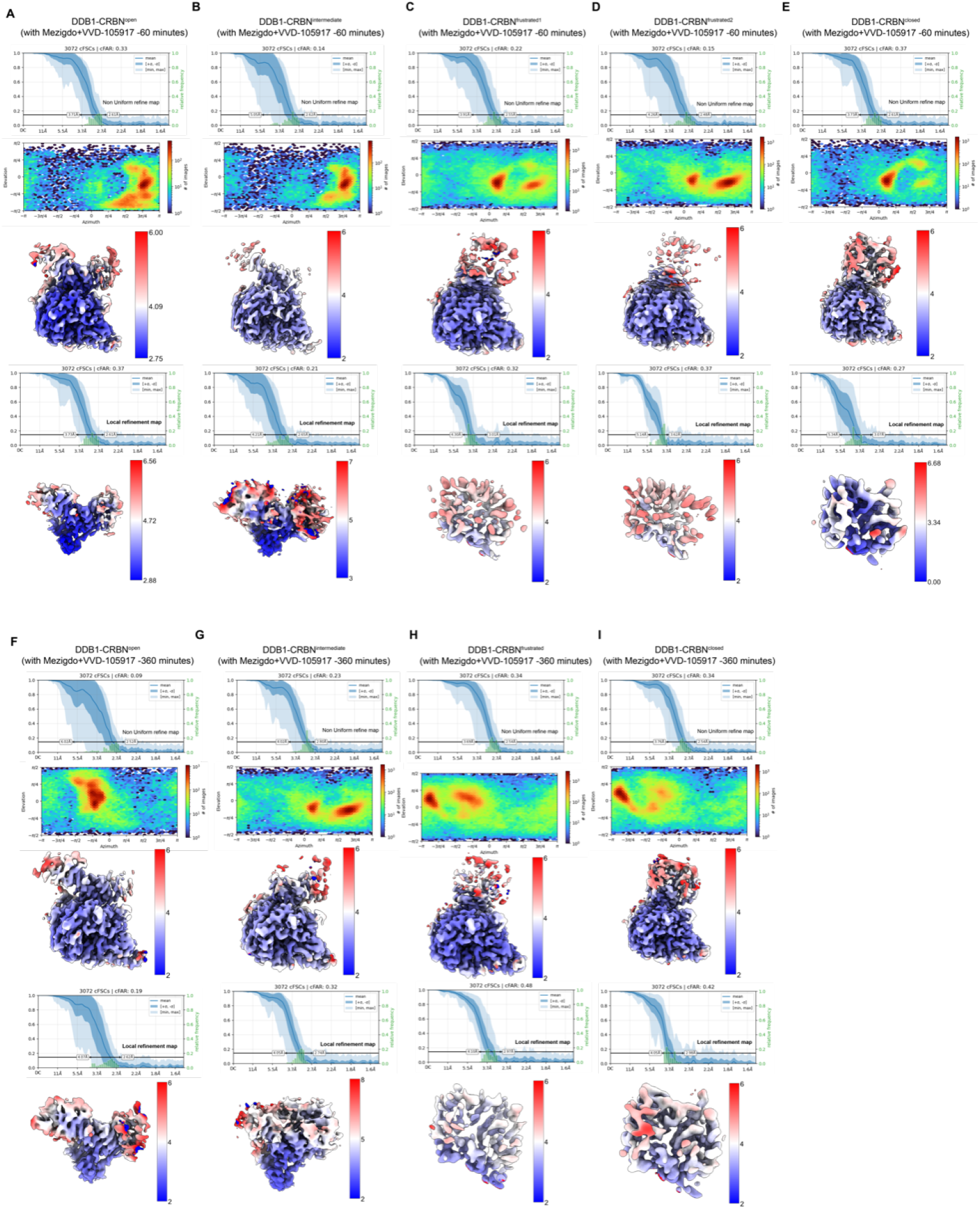
Resolution, angular distribution, and local resolution assessment of cryo-EM reconstructions. **(a-i)** Cryo-EM data quality metrics for each reconstruction. Each panel contains, from top to bottom, a conical Fourier shell correlation (cFSC) curve, with overlaid histogram of directional resolutions, an angular distribution plot, the full reconstruction colored by local resolution; a cFSC curve for masked local refinement of the CRBN domain, and the locally refined CRBN domain colored by local resolution. The reconstructions are as follows: **a-e**, DDB1^ΔBPB^/CRBN^open^ (a), DDB1^ΔBPB^/CRBN^intermediate^ (b), DDB1^ΔBPB^/CRBN^frustrated1^ (c), DDB1^ΔBPB^/CRBN^frustrated2^ (d), and DDB1^ΔBPB^/CRBN^closed^ (e) after incubation with mezigdomide for 10 minutes followed by VVD-105917 for 60 minutes. **f-i**, DDB1^ΔBPB^/CRBN^open^ (f), DDB1^ΔBPB^/CRBN^intermediate^ (g), DDB1^ΔBPB^/CRBN^frustrated^ (h), and DDB1^ΔBPB^/CRBN^closed^ (i) after incubation with mezigdomide for 60 minutes followed by VVD-105917 for 360 minutes.

**Supplementary Table 1.**
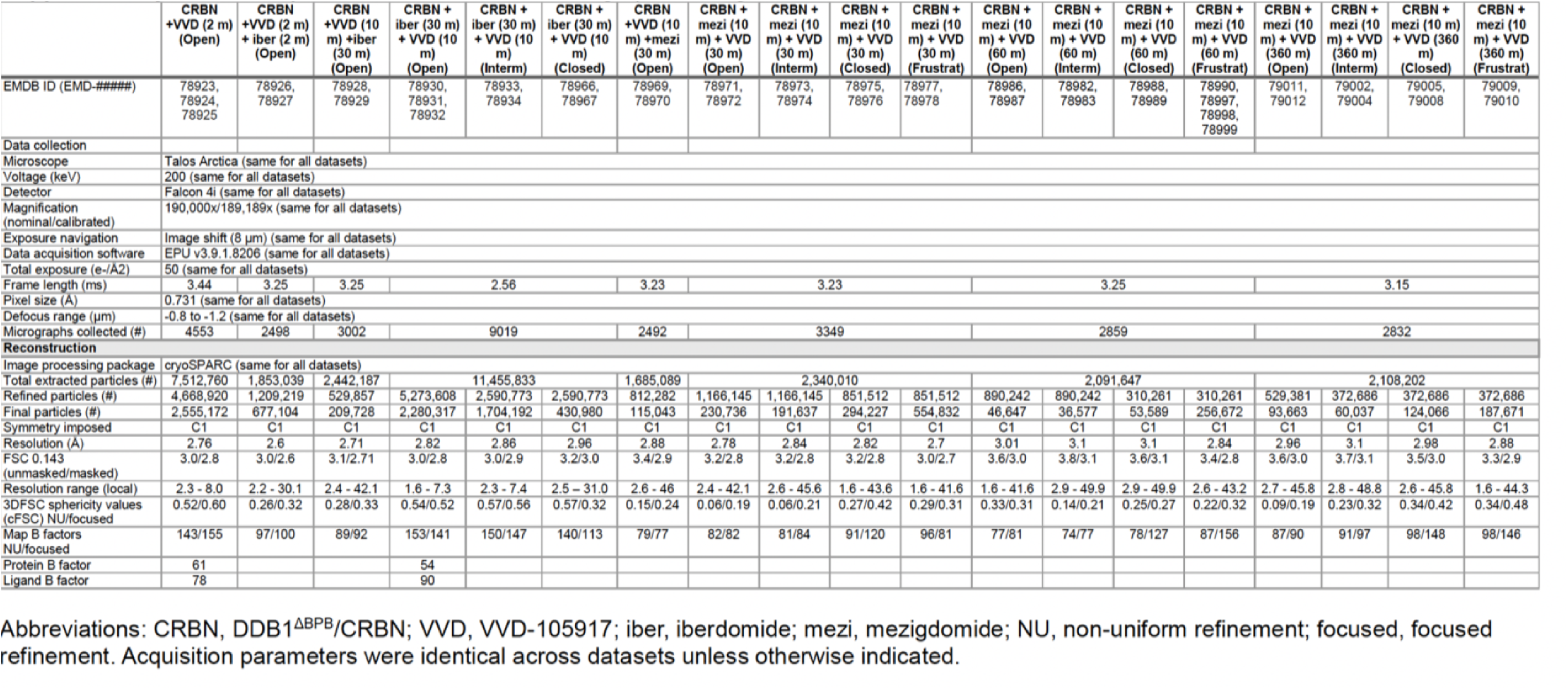
Cryo-EM data collection and image-processing statistics.

**Supplementary Table 2.** Atomic modeling, refinement, and validation statistics.

|  | DDB1<sup>ΔBPB</sup>/CRBN +<br>VVD-105917 (2 m) | DDB1<sup>ΔBPB</sup>/CRBN +<br>iberdomide (30 m) + VVD-105917 (10 m) |
| --- | --- | --- |
| **PDB ID** | 38MG | 38MJ |
| **Model composition** |  |  |
| Protein residues | 1183 | 1145 |
| Ligands | 2 | 3 |
| **Model refinement** |  |  |
| Refinement package | Phenix v2.0orc1-5617 | Phenix v2.0orc1-5617 |
| CC (Volume/mask) | 0.86/0.85 | 0.88/0.88 |
| RMS deviations |  |  |
| Bond lengths (Å) | 0.004 (0) | 0.004 (0) |
| Bond angles (°) | 0.583 (2) | 0.706 (3) |
| Validation |  |  |
| Map to model FSC 0.5 | 1.7 | 2.1 |
| Ramachandran (%) |  |  |
| Outliers | 0 | 0 |
| Allowed | 4.8 | 4.92 |
| Favored | 95.2 | 95.08 |
| MolProbity score | 1.81 | 1.97 |
| Poor rotamers (%) | 2.28 | 2.44 |
| Clash score (all atoms) | 4.26 | 6.11 |
| C-beta deviations | 0 | 0 |
| CaBLAM outliers (%) | 2.87 | 2.38 |
| Q score | 0.81 | 0.75 |
| EMRinger score | 4.19 | 3.46 |

## Notes

### Competing Interest Statement

B.E.N., M.P.P., and G.M.S. are current employees of Vividion Therapeutics. K.A.B. was an employee of Vividion Therapeutics when the research was conducted. G.C.L. is a current employee of Bristol Myers Squibb. The remaining authors declare no competing interests.

